# Modelling the Perception of Colour Patterns in Vertebrates with HMAX

**DOI:** 10.1101/552307

**Authors:** Julien P. Renoult, Bastien Guyl, Tamra C. Mendelson, Alice Percher, Jérôme Dorignac, Fredäric Geniet, Molino Franęois

## Abstract

1. In order to study colour signals as animals perceive them, visual ecologists usually rely on models of colour vision that do not consider patterns–the spatial arrangement of features within a signal.
2. HMAX describes a family of models that are used to study pattern perception in human vision research, and which have inspired many artificial intelligence algorithms. In this article, we highlight that the sensory and brain mechanisms modelled in HMAX are widespread, occurring in most if not all vertebrates, thus offering HMAX models a wide range of applications in visual ecology.
3. We begin with a short description of the neural mechanisms of pattern perception in vertebrates, emphasizing similarities in processes across species. Then, we provide a detailed description of HMAX, highlighting how the model is linked to biological vision. We further present *sparse-HMAX,* an extension of HMAX that includes a sparse coding scheme, in order to make the model even more biologically realistic and to provide a tool for estimating efficiency in information processing. In an illustrative analysis, we then show that HMAX performs better than two other reference methods (manually-positioned landmarks and the SURF algorithm) for estimating similarities between faces in a nonhuman primate species.
4. This manuscript is accompanied with MATLAB codes of an efficient implementation of HMAX and sparse-HMAX that can be further flexibly parameterized to model non-human colour vision, with the goal to encourage visual ecologists to adopt tools from computer vision and computational neuroscience.

## Introduction

Understanding the evolution and the ecological significance of communicative traits requires studying these traits in the eyes of beholders (Endler *et al.* 2005). In visual communication, colour spaces-which model perceived differences between colours-have thus become very popular among visual ecologists who study socio-sexual communication, camouflage, mimicry and plant-animal interactions (Renoult, Kelber & Schaefer 2017). In colour spaces, the design of visual stimuli is usually studied as a collection of isolated plain colours, without considering the influence of their spatial arrangement on perception. However, the effectiveness of a communication system strongly depends on how colour patches, lines or dots are arranged spatially to form colour patterns. For example, the diurnal hawkmoth *Macroglossum stellatarum* innately prefers radial blue and white patterns to ring patterns with the same colours (Kelber 2002). Modelling the perception of colour patterns is thus a necessary step toward a better understanding of natural communication systems.

In this article, we highlight the benefits of the HMAX family of models for analysing patterned colour stimuli, as vertebrates perceive them. HMAX was originally developed by computational neuroscientists to model information processing in the ventral stream of the visual pathway, that is, the brain area involved in shape and colour perception in humans (Serre & Riesenhuber 2004). Yet, due to the generality of the hierarchical mechanisms involved in this family of models, the sensory and brain processes modelled in HMAX are certainly widespread, occurring in most if not all vertebrate taxa, thus offering HMAX a wide range of applications in visual ecology.

We begin with a short description of the neural mechanisms of pattern perception in vertebrates, emphasizing similarities in processes across species. Then, we provide a detailed description of HMAX, highlighting how it is connected to biological vision. We further present sparse-HMAX, an extension of HMAX that includes a sparse coding scheme. Sparse coding describes the strategy of neural systems to minimize the number of neurons activated simultaneously (Olshausen & Field 2004). By adding sparse coding to HMAX, we aim both to develop an even more biologically realistic model of perception of colour patterns, and to provide a framework for estimating efficiency in information processing (Renoult & Mendelson 2019). A fast and easily customizable version of HMAX that can be used to model the perception of colour patterns in most vertebrates and our sparse-HMAX algorithm are available at: https://github.com/EEVCOM-Montpellier/HMAX. In the last part of this article, we apply HMAX to estimate similarities between faces in a nonhuman primate species as an example application.

## Perception of colour patterns in vertebrates

Despite structural differences in how vertebrates perceive colour patterns, a number of general principles governing the processing of visual information are shared across species. In this section, we review four of these principles: the hierarchical processing of information, the tuning of neurons to stimulus features, the sparse encoding of information, and the opponent processing of colour information.

### Hierarchical processing of visual information

The perception of colour pattern is one step of the whole vision process that ultimately leads to recognition. In mammals, vision starts with the stimulation of retinal photoreceptors that convert the light arising from a stimulus into electro-chemical signals. These signals are then conducted through retinal ganglion cells to reach the lateral geniculate nucleus (LGN), a relay centre that connects the retina to the primary visual cortex (V1). Signals continue to flow bottom-up through V2 and V4, and then through the inferior temporal cortex (IT), which feeds the prefontal cortex that connects perception to memory and action (Felleman & Van Essen 1991). Areas V3 and V5 are involved mainly in motion vision (Zeki *et al.* 1991).

As signals flow from photoreceptors up to IT, the information extracted becomes increasingly complex (Mély & Serre 2017). At the receptor level, light contrasts are recorded locally without any information about their spatial organization. In V1, neurons become sensitive to short and oriented line segments (Tootell *et al.* 1988). Basic shapes such as curved lines (i.e. combinations of oriented line segments) are mainly processed in V4. More complex shapes representing entire objects (i.e. combinations of curved lines; e.g., a lion, a house or a face) are processed in IT and in subsequent specialised areas (e.g., the fusiform face area for faces). In addition, throughout the visual pathway neurons are increasingly invariant to orientation, scale, position and lighting conditions. Neurons in IT thus fire in response to specific items yet they are insensitive to how tilted, distant, centred in the field of view and shaded these items are (Mély & Serre 2017). How the visual system achieves the dual increase in sensitivity and invariance has been a central question of vision science and is still one of the most active research topics in computer vision (e.g., Anselmi *et al.* 2016).

In their seminal article, Hubel and Wiesel (1962) proposed a physiological model of V1 that copes with the complexity-invariance problem. The model assumes a feedforward, hierarchical flow of information within V1 that involves two different types of neurons: the simple cells and the complex cells. Simple cells pool afferents (LGN cells with circular receptive fields; RFs hereafter) sampled along oriented line segments. Complex cells pool inputs from several spatially contiguous simple cells with different orientations, phases, positions or scales. Consequently, one complex cell keeps the complex selectivity of its afferent simple cells but it is tolerant to local variation in stimulus orientation, phase, position or scaling. This hierarchical model has been extended to V2, V4 and IT, were neurons equivalent to simple and complex cells have been discovered (Soto & Wasserman 2012). Information thus flows through the visual pathway alternating between simple cells, with increasingly complex selectivity, and complex cells, with increasingly large tolerance to geometrical transformations. Furthermore, because both types of cells pool information from multiple afferents, the neurons’ RFs become increasingly larger as signals flow up (Smith *et al.* 2001).

The hierarchical processing of visual information is a general principle found in other, non-mammalian vertebrates. Most neurons in the tecto-isthmic system–the functionally analogous structure to the visual cortex in non-mammals–are selective to orientation (e.g., in fish see Ben-Tov *et al.* 2013). In birds, it was found that the elongated RF of isthmic cells is generated by pooling afferents from aligned tectal cells with circular RFs, and that this elongated RF underlies the orientation selectivity of these cells (Li, Xiao & Wang 2006). In fishes, the laminar organisation of orientation-selective inputs coming from the retina suggests that different layers in the tectum may be dedicated to processing specific visual features (Abbas & Meyer 2014). RF size has been shown to increase along the bird visual pathway too (Engelage & Bischof 1996). Overall, the available behavioural and neurophysiological data indicate that, although they use ontogenetically different structures, a similar hierarchical and feedforward processing occurs in the tecto-isthmic system and in the visual cortex (Soto & Wasserman 2012).

Besides feedforward projections, the visual pathway of vertebrate vision also involves horizontal and feedback neural projections (Treue 2003). Yet, because of the short response latency of IT’s neurons to visual stimuli (~100 ms), and the ability of primates to recognize objects in a very short time (~150 ms; Thorpe, Fize & Marlot 1996), it is generally recognized that core object representation, and *a fortiori* pattern perception, are essentially feedforward mechanisms (VanRullen & Koch 2003; DiCarlo, Zoccolan & Rust 2012). This is why the simple hierarchical model of Hubel and Wiesel has been consistently efficient in explaining and predicting empirical results in neuroscience over the years (Ferster & Miller 2000; Reid & Usrey 2004).

### The tuning of neuronal selectivity

For visual ecologists, an important question is whether the selectivity of cortical/tecto-isthmic neurons adapts to the environment, and whether this adaptation is determined developmentally or evolutionarily. In contrast to many studies supporting a spectral tuning of photoreceptors to lighting conditions (listed in Cummings & Endler 2018), analyses comparing the shape and orientation selectivity of neurons between species inhabiting contrasting visual environments are still lacking. Nevertheless, a few studies with model species reveal interesting relationships between orientation selectivity and environmental stimuli. One study in kittens showed that, at eye opening, a proportion of cells in V1 show the orientation selectivity typical of a mature visual cortex (Sengpiel & Kind 2002). Moreover, the visual cortex of kittens reared in a striped environment responded to all orientations, even those never seen by the animal, even though twice as much cortical area was devoted to the experienced orientation (Sengpiel, Stawinski & Bonhoeffer 1999). For more complex shapes, and thus in higher levels of information processing, there is also evidence that some stimuli are innately categorized (e.g., faces in primates: Johnson *et al.* 1991); nevertheless it is generally accepted that complex shape selectivity is mostly tuned by learning (e.g., Freedman *et al.* 2005). Overall, it appears that shape selectivity is innate, but that it can be retuned to environmental stimuli through learning in some neurons; moreover, the proportion of neurons that can be retuned seems to increase in higher levels of the visual pathway.

### Sparse coding

At a given level of information processing, signals from individual neurons are combined into a neural code, i.e. a neural representation of a visual stimulus at this particular level. Yet the high metabolic cost of neuronal activation (in the human visual system, it accounts for 2.5 to 3.5 % of a resting body’s overall energy requirements; Attwell & Laughlin 2001) imposes constraints onto the neural code (Graham & Field 2006). Olshausen and his colleagues analysed the importance of this constraint by training artificial neurons to encode images of natural scenes as efficiently and precisely as possible (Olshausen & Field 1996; Olshausen & Field 1997). To match the selectivity of these artificial neurons to that of real neurons measured in mammalian V1, the authors had to implement a sparseness criterion in the neural code. In this context, sparseness can describe both the fact that only a small fraction of neurons are active at any time (population sparseness), and that individual neurons activate shortly and rarely (temporal sparseness). Sparse coding is metabolically effective because frequently firing a few generalist neurons is far more costly than maintaining a large population of highly selective yet sparsely activated neurons (Lennie 2003; Olshausen & Field 2004). Experimentally, following its discovery in V1, sparse coding has been demonstrated at all levels of perception, from the retina (Pitkow & Meister 2012) to V4 (Carlson *et al.* 2011) and IT (Brincat & Connor 2004). Sparse coding is also ubiquitous in other sensory modalities and has been found in all organisms investigated so far, including invertebrates (e.g.; Hromádka, DeWeese & Zador 2008).

### Opponent colour coding

The studies described previously have investigated the selectivity of neurons to luminance (i.e. along a greyscale) contrasts; yet most vertebrates additionally use colour as a source of information. It is well established that opponent coding is crucial for modelling colour perception: compared to analysing raw excitation values, analysing *differences* in photoreceptor excitations dramatically improves the fit between predicted and actual perceived differences between colours (Renoult, Kelber & Schaefer 2017). In Old World primates, colour opponency is achieved by combining outputs of the three S, M and L photoreceptors (standing for short, medium and long wavelength, respectively) into two opponent channels: the red-green and the blue-yellow channels computed as L-M and S-(M+L), respectively. However, the predictive power of these two opponent channels strongly depends on the size and shape of the stimulus (Derrington, Krauskopf & Lennie 1984). Although it has been traditionally assumed that luminance and colour are processed separately in the visual system of vertebrates, and that shape perception is mediated by the luminance channel, evidence has accumulated that colour and shape are inextricably linked (for a review, see Shapley & Hawken 2011).

The neurons that compute the L-M and S-(M+L) signals represent only one of two categories of opponent cells found in the vertebrate visual system, which have been named single-opponent (SO) cells (Shapley & Hawken 2011). SO cells compute an averaged difference in photoreceptor excitation locally, and thus they are mostly activated by full-field stimuli. SO cells are useful for processing surface information and are almost not selective to orientation. Another category of colour-selective cells receives inputs from SO cells. These so-called double-opponent (DO) cells compute an averaged difference in colour (L-M or S-(M+L)) signals between different regions of the visual field (Shapley & Hawken 2011). DO cells located in the retina and LGN have a circular RF and are thus unselective to orientation. In V1, however, many DO cells have an oriented RF (Johnson, Hawken & Shapley 2001). Oriented DO cells are useful for processing coloured line segments and thus are very important for the perception of colour patterns. Furthermore, DO cells are selective to both colour and luminance contrasts, which argues against the idea of a strong segregation of colour and luminance processing beyond the LGN (Gegenfurtner 2003). Besides primates, SO and DO cells have been found in other vertebrates including fishes (Daw 1968) and birds (Frost, Scilley & Wong 1981).

In summary, the perception of colour patterns in vertebrates relies on general principles that are likely shared among most species: a hierarchical processing of signals, alternating shape-selective and spatial-pooling neurons, the adaptation of shape-selective neurons, sparse coding and a simple/double opponent colour coding scheme.

## HMAX and spαrse-HMAX

For more than five decades, researchers from cognitive sciences, computer vision and artificial intelligence have proposed mathematical models of colour pattern perception that account for the aforementioned principles (reviewed in Poggio & Serre 2013; Serre 2013). HMAX (Riesenhuber & Poggio 1999; Serre *et al.* 2005) is one of the most popular families of models of visual processing in humans. The original HMAX has been improved in multiple ways; the version presented here is generalizable to non-human vertebrates and can further account for sparse coding.

### HMAX

HMAX is a hierarchical neural network starting with a scale image *I* as an input layer and alternating *S* layers that perform feature mapping and *C* layers that perform feature pooling. *S* and *C* stand for “Simple” and “Complex”, in reference to the simple and complex cells described previously. Feature mapping is achieved by convoluting different shape-selective filters (or artificial neurons) with *I* in the first layer, or with maps produced by the preceding layer. Feature pooling increases the invariance of feature maps to geometrical transformations. Classical versions of HMAX include four layers in addition to the input layer: *S*1, *C*1, *S*2 and *C*2 (Figure 1a). The output of C2 is a compact code describing the input image, typically a vector of a thousand values.

**Figure 1.**
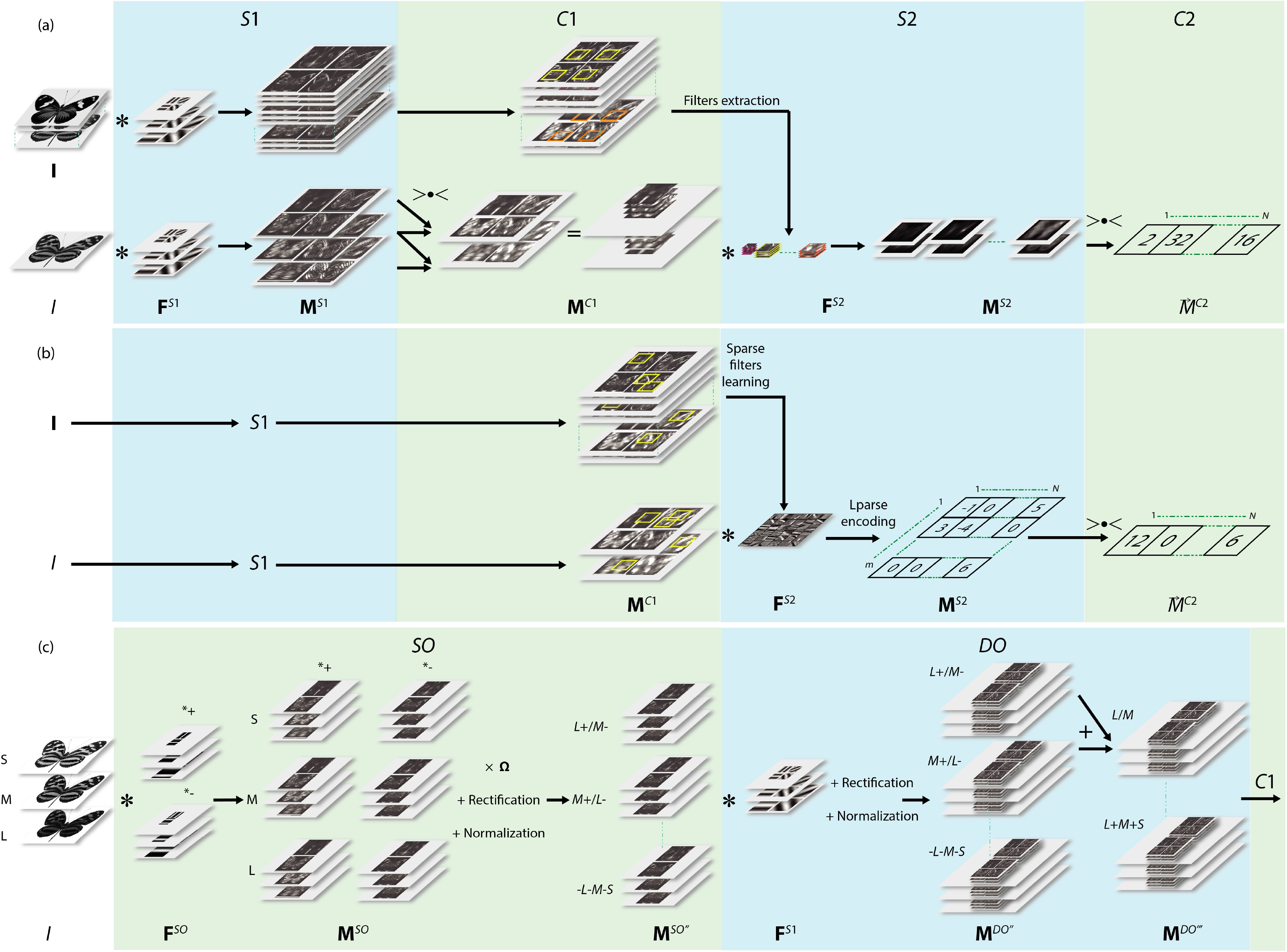
Overview of HMAX models. (a) Classical HMAX for scale images with filter learning (upper row) and stimulus encoding (lower row). (b) sparse-HMAX for scale images with filter learning (upper row) and stimulus encoding (lower row). (c) Simple opponent (SO) and double-opponent (DO) processes in HMAX for colour images (here, describing photoreceptor excitation maps of a trichromat species).

*Layer S1*.- In *S*1, feature-selective neurons are represented by a set **F**^*S*1^ of Gabor filters 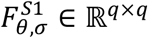 defined by orientation *θ* ∈ ℝ^*t*^ (t different values), scale *σ* ∈ ℝ^*sc*^ and filter size 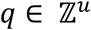. Gabor filters have been previously shown to accurately model the selectivity both of simple cells of V1 in mammals (Jones & Palmer 1987), and also (most likely) of tecto-isthmic neurons with an elongated RF in other vertebrates. Defining *x_g_*=1,…,*q* and *y_g_*=1,…,*q*, a Gabor filter 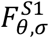 is described as

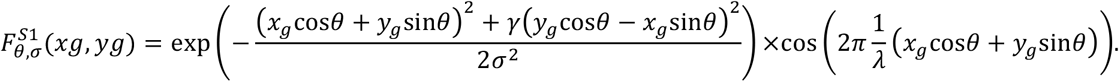

with *γ* the aspect ratio of the filter and *λ* the wavelength parameter. Serre and Riesenhuber (2004) found that *λ* had limited effect on tuning filter selectivity and thus kept it constant. These authors further proposed to approximate *σ* and *λ* from *q* such that

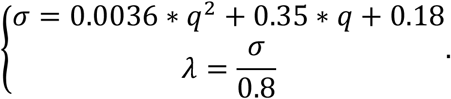

The number of different scales (*sc*) is thus taken as identical to the number of different filter sizes (*u*). **F**^*S*1^ is determined by the set of *SC/u* and *t* values for *σ/q* and *θ* respectively. Those should vary between species and, as discussed previously with the example of kittens reared in a striped environment, with experience (for parameter values that best fit to neurophysiological data in mammals, see Serre & Riesenhuber 2004). Having built the filter set **F**^*S*1^, following Theriault, Thome & Cord (2013), each filter is then convolved with *I* to generate *txsc* feature maps 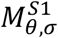 such that

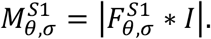

*Layer C1.– C1* pools values of 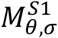 to produce 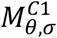 maps with a decreased resolution but an increased tolerance of features to shift and size (but not to orientation). Among the different pooling functions, max pooling (keeping the maximum value only) works best in generating invariant features (Scherer, Müller & Behnke 2010). For each of the *t* values of the orientation *θ,* a max filter *F*^*c*1^ ∈ ℝ^*rxr*^ (with *r* ∝ *q*, see Theriault, Thome & Cord 2013) is applied simultaneously to neighbourhood values within a given 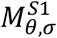 feature map (thus gaining position invariance), and to pairs of maps with two consecutive scales (thus gaining scale invariance). The max filter is moved in steps of 1 between scales, and around a given feature map in steps of magnitude r. Note that due to the pairwise pooling in scale, in *C*1 there is only *SC*-1 remaining values for *σ.*

*Layer S2.-* V4/IT/isthmic neurons are modelled in *S*2 by a set **F**^*S*2^ of filters, which are selective to more complex features (curved lines, shapes) compared to 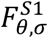 (segment lines; Figure 2). Motivated by the biological evidence that shape-selectivity is largely tuned by adaptation or experience to environmental stimulation in higher levels of the visual pathway, **F**^*S*2^ is generated in an initial feature learning stage using a set of training images. This stage includes four steps. First, 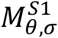 and then 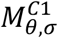 are generated for each learning image. Second, the coordinates (x,y) of a filter centre are randomly drawn in 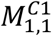. Third, one filter size *q*′ is randomly drawn, with the constraint that the different possible values of *q*′ are all equally represented in **F**^*S*2^.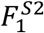 is then extracted from 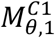 around (x,y) at the first scale *σ* =1 and for all orientations. Steps two to four are repeated to generate *N* filters 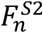; typically *N* = 10^3^. Note that each 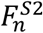 is thus constructed as a 3D filter: ℝ^*q*′x*q*′x*t*^. In contrast to *S*1, the learning stage of *S*2 allows shape-selective filters to adapt to dominant shapes within training images, which can be the same as the target stimulus or other images depicting, e.g., environmental scenes.

**Figure 2.**
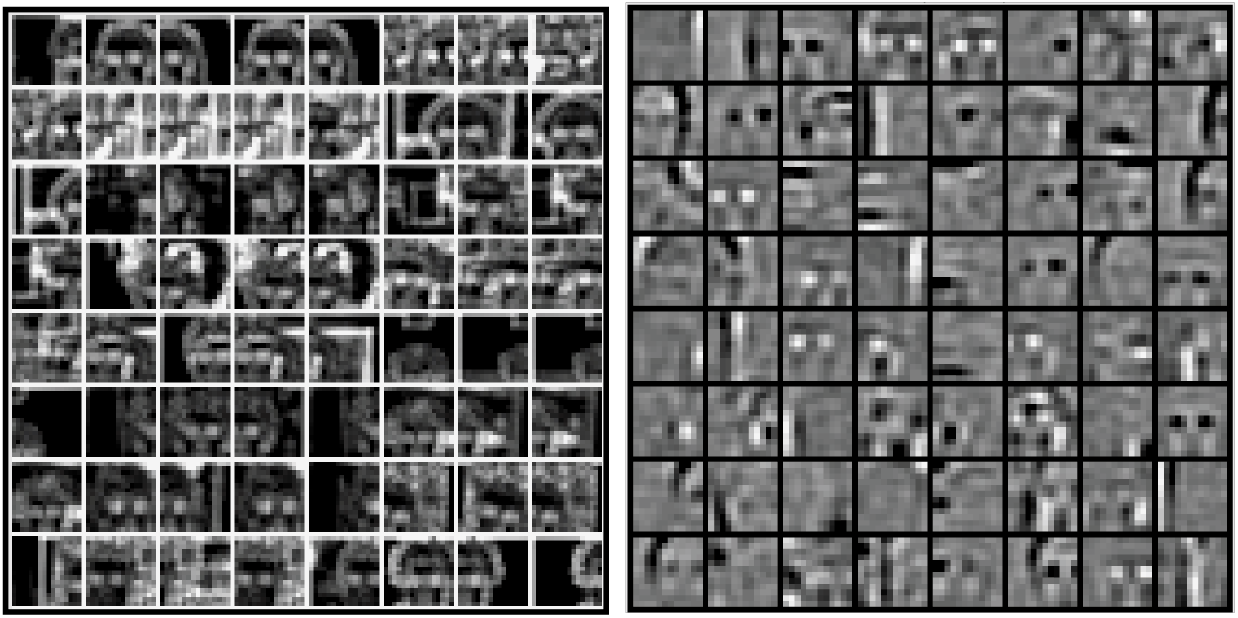
Filters in S2 (**F**^S2^) learned on the human face category of Calthech101 image dataset with classical HMAX (left) and *sparse*-HMAX (right). Filters in S2 model neurons selective to complex features and with large receptive fields (here, regions of faces).

The output of *S*2, i.e., feature maps 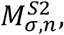, are eventually built by convoluting 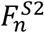 with 3D stacks of 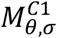,

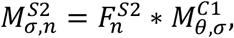

using the radial basis function described in (Mutch & Lowe 2008) that calculates, at every position (*x,y*) of the map, the response *Re* to the filter of a patch *P* from 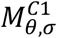 as

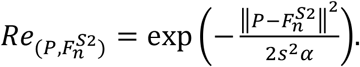

Mutch & Lowe (2008) proposed to set *s* to unit value and the normalization factor *α* to (*q*/4)^2^, with *q* set to the smallest possible value. The convolution operation eventually generates *N*x(*sc* – 1) different maps.

*Layer C2.– C2* pools values of 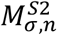 feature maps to gain global invariance to position and scale. For each *n* max pooling is applied across all positions and scales to produce a vector 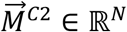, the output of HMAX.

### sparse-HMAX

Sparse coding can be added to HMAX at two different stages: during filter learning (e.g., Hu *et al.* 2014) or during stimulus encoding. In computer vision these stages are clearly distinct: it is possible to sparsely encode a stimulus with Gabor filters, or to encode a stimulus with a simple linear convolution with filters learned by applying a sparseness constraint. In *sparse*-HMAX, the user can choose to impose sparseness during filter learning, during stimulus encoding in layer *S*2 (Figure 1b), or at both steps. In filter learning, *m* square patches are first extracted from 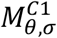 of training images. Each patch is then transformed into a vector 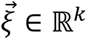, and all 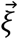 are h-concatenated into a matrix **X** ∈ ℝ^*k x m*^. **X** is then unity-based normalized and zero-centred. Learning sparse filters requires solving the following optimisation problem:

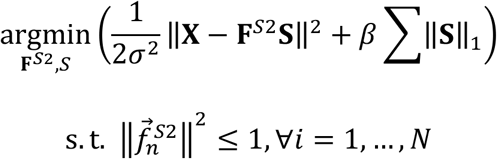

where **F**^*S*2^ ∈ ℝ^*k x N*^ is a matrix of *N* vectorised filters 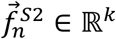 (the set of filters that are learned) and **S** ∈ ℝ^*N x m*^ a matrix of filter weights. The left part of the problem, called the reconstruction error, is a least square minimisation of the difference between observed patches and patches predicted by a linear combination of filters. The right part of the problem, called penalty function, imposes sparseness to **S**. Here the penalty function is a *ℓ*_1_-norm regularisation, which is notoriously good at generating sparse weights while being robust to irrelevant features (Ng 2004). The regularisation parameter *β* determines the relative importance given to maximising either the reconstruction accuracy or the sparseness. To prevent **F**^*S*2^ from having arbitrarily large values that would lead to arbitrarily small values in **S**, its columns 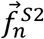 are constrained to have their *ℓ*_2_-norm less than or equal to one. Since the problem in convex in **F**^*S*2^ (while holding **S** fixed) and convex in **S** (while holding **F**^*S*2^ fixed) but not convex in the two simultaneously, it is necessary to iteratively and alternatively minimize with respect to **F**^*S*2^ or **S** while holding the other element fixed. Theoretically, this iterative optimisation could be achieved using a metaheuristic evolutionary algorithm using operators such as reproduction, mutation and recombination, in order to simulate an evolutionary adaptation of the neuronal selectivity to environmental features. In practice, however, the time needed to solve the problem would be unrealistically long, and several more efficient algorithms have been developed. In our implementation of sparse-HMAX, we used the *Fast Sparse Coding* algorithm (Lee *et al.* 2007), which derives and solves the Lagrange dual to learn the filters and applies a feature-sign search to learn the weights (for details, see Lee *et al.* 2007).

To sparsely encode a target stimulus, as in training, a matrix **X** of *m* square patches is extracted in a similar way from 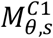. maps of the target image *I*, then normalised and centred, and eventually the feature-sign search algorithm is used to learn **S** using **F**^*S*2^, the set of filters learned previously. In sparse-HMAX, the output of *S*2 is thus **S**, which is equivalent to a single feature map indicating the activation of each filter for all patches (and thus for all scales and orientations in C1). The output 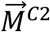 of *sparse-HMAX* is eventually given by

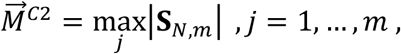

that is, by extracting the maximal activation of each filter over all patches.

### Modelling colour opponency

Classical models of HMAX have been developed to process visual stimuli in the luminance channel only. However, Zhang, Barhomi & Serre (2012) recently developed coloured feature maps that model single- and double-opponent cells in primate V1, and proposed an implementation of these maps into HMAX (see also Mély & Serre 2017). In the following, we present the *SO-DO* feature maps of Zhang, Barhomi & Serre (2012) generalized to any opponent function and any number of photoreceptor types used in colour vision (e.g., to dichromatic mammals or to tetrachromatic birds; Figure 1c). Let set *I* ∈ *ℝ*^*hxwxnpe*^ the input matrix of photoreceptor excitation maps *I_b_* ∈ *M^hxw^*, with *b* ∈ ℤ^*npe*^. In Old World primates, *npe* = 3 with *b* ∈ {*L, M, S*}, which is often approximated by {*R, G, B*}, the blue, green and red channels of a colour image. Moreover, in some species like primates and fishes, the luminance channel is given by summing two or more *I_b_* (see *Layer SO*). In other species, e.g., in birds, a specific set of photoreceptors (the double cones) feeds the luminance channel. In this case, the photoreceptor excitation map of the luminance channel should be included as a specific *I_b_* within *I*.

*Layer SO.-* The spatio-chromatic selectivity of *SO* cells is modelled using Gabor filters as in the non-colour HMAX. However, here one Gabor filter is decomposed into two filters: one keeping the positive part of the filter values only, to model an excitatory cell (other values are set to zero), the other one keeping the negative values to model an inhibitory cell. *SO* filters are thus defined by 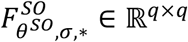, with *θ*^*so*^ ∈ ℝ^t^SO^^, *σ* ∈ ℝ^*sc*^ and * ∈ {ℝ^-^, ℝ^+^} indicating the filter polarity (excitatory or inhibitory). Because *SO* cells are only weakly selective to orientation, Zhang, Barhomi & Serre (2012) set *t^SO^* = 2 for humans, which likely also applies to other vertebrates. Having built the filter set **F**^*SO*^, each filter is then convolved with each photoreceptor excitation map to generate a set **M**^*SO*^ of *t*^*so*^xscx2xnpe feature maps 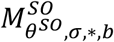 such that

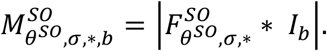

The set of single opponent and luminance maps **M**^*SO*^’ containing *t*^*so*^*xscx2xnch* maps 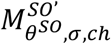 with *ch* ℝ ℤ^*nch*^ are then obtained by linearly combining 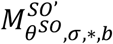 such that

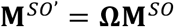

with Ω ∈ ℝ^*npexnch*^ is a matrix of weights in which columns define the single opponent and luminance functions. For example, in Old World primates,

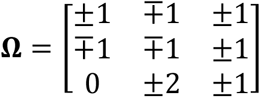

thereby producing, for each orientation *θ^SO^* and scale *σ*, four opponent colour channels: *L*+/*M*- (red excitatory and green inhibitory), *M*+/*L*-, *S*+/(*M*+*L*)-, (*M*+*L*)+/*S*-; and two luminance channels: *L*+*M*+*S*,-*L*-*M*-*S*.

Each 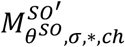 map is then rectified by half-squaring to maintain positive firing rate (Heeger 1992; Zhang, Barhomi & Serre 2012):

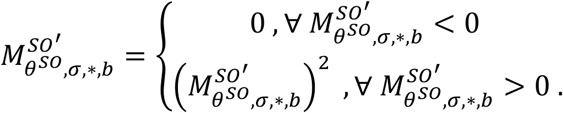

The last step of layer *SO* is a divisive normalisation that provides tolerance to small light intensity scaling:

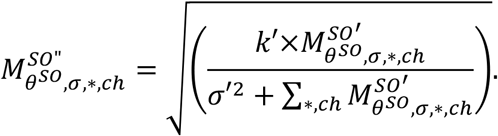

Note that in this layer normalisation is performed over maps of all (excitatory and inhibitory) channels at a given *θ^SO^* and *σ*. Based on neurophysiological data from macaque, Zhang, Barhomi & Serre (2012) fixed the scale *k*′ and semi-saturation *σ*′ parameters to 1 and 0.225, respectively.

*Layer* DO.-Each map of *SO* is first convolved with the classical Gabor filters 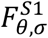 (thus not dissociating excitatory and inhibitory subunits), which produces *t^so^xscx2xnchxt* maps 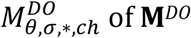:

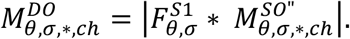

Each 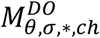 map is then rectified by half-squaring as in *SO*, to produce 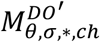. A normalisation is further applied, but over orientations (instead of channels in *SO*):

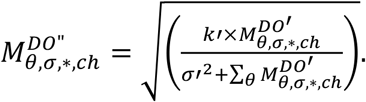

Last, feature maps 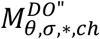 with complementary channels (excitatory and inhibitory) are summed in order to make *DO* cells insensitive to figure-ground reversal:

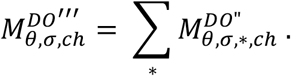

From there, all of the *t*x*sc*x*nch* maps of **M**^*DO*^”’ are processed individually by *C*1, *S*2 and *C*2, and all 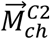 are concatenated to produce 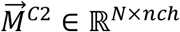, the final output of the HMAX and *sparse*-HMAX.

### Example: estimating facial (dis) similarity in mandrills

HMAX can be used to estimate the perceived resemblance between entire phenotypes. To illustrate this, we computed the distance between vectors *C*2 encoding faces of mandrills *(Mandrillus sphinx*) and compared the results with a method based on manually-positioned landmarks that is widely used in primatology (Dal Martello & Maloney 2006; Bower, Suomi & Paukner 2012). In addition, we compared HMAX to another computer vision approach based on the number of shared, automatically detected features, which has been previously used in evolutionary ecology (Stoddard, Kilner & Town 2014; SURF method hereafter). We analysed 100 pictures of mandrill faces depicting 75 different individuals. Twenty-six individuals were represented by more than one portrait, taken between 6 and 36 months intervals. For each of the four methods (landmark, SURF, classic HMAX and *sparse*-HMAX), we estimated the similarity between all pairs of portraits (for details, see Supporting Information), we ordered pairs by ascending value of similarity and summed the rank of pairs corresponding to different pictures of the same individuals. The higher the rank sum, the best the method to assign high similarity to same-individual portrait pictures. For all three methods, the rank sum was higher than expected by chance (95% limit of a null distribution), indicating that all methods recognized that same-individual pictures were more similar than different-individual pictures (Table 2). The landmark and SURF methods reached similar performance, but both methods were surpassed by HMAX. Performance of sparse-HMAX was lower than that of HMAX but higher than that of landmark and SURF methods.

**Table 2.** Results of the facial similarity analysis. The standardized (Sd) rank sum corresponds to the rank sum divided by that of an ideal observer who would give the highest rank to all same-individual pairs.

| Method | Rank Sum ($\times 10^3$) | Sd rank sum | *p*-value |
| --- | --- | --- | --- |
| 95% limit | 84.4 | 0.59 | 0.05 |
| Landmark | 99.5 | 0.70 | 1e-4 |
| SURF | 100.2 | 0.70 | 4e-5 |
| HMAX | 111.3 | 0.78 | <1e-6 |
| *sparse*-HMAX | 104.9 | 0.73 | <1e-6 |
| Ideal observer | 143.1 | 1 | <1e-6 |

## Discussion

With the increasing need to study colour patterns as animals perceive them, HMAX offers a useful framework applicable to a wide array of vertebrate species. We provide a flexible and fast (See performance tests in Supporting Information) implementation of HMAX that can apply to RGB images, but also to the photoreceptor excitation spaces of other vertebrates. Furthermore, photoreceptor excitations can be combined into any opponent coding scheme. Knowing the visual acuity of a studied species (for a review, see Caves, Brandley & Johnsen 2018), it should be possible to further set the scale parameter of Gabor filters to realistically map the visual resolution of details in colour patterns. Even without specific knowledge about the visual system of a species, HMAX would still be a valuable tool to analyse the robustness of results (e.g., measures of phenotypic similarity) to deviations from hypotheses about the visual processes; e.g., the uniform distribution of orientation-selective neurons, or the adaptation of neurons in early cortical/tecto-isthmic areas to features of the environment.

HMAX also allows analysing communication signals independently of their rotation, distance and illumination. For visual ecology, this is a critical advantage over the classical landmark-based methods because it allows working with non-standardized images, e.g., pictures collected on the World Wide Web, thereby opening up the possibility to analyse very large number of images or rare species represented by low-quality photographs only. Furthermore, automatic analysis of features will save considerable amount of time compared to positioning landmarks manually.

Over the last few years, computer scientists have gradually shifted away from HMAX models to favour artificial neural networks with deeper architectures (i.e. deep convolutional neural networks; ConvNets), reaching performances in object and individual recognition that match and even surpass those of humans (LeCun, Bengio & Hinton 2015). ConvNets have also recently gained the interest of visual scientists with their ability to predict the selectivity of biological neurons in different brain areas (Kriegeskorte 2015). However, HMAX models have two main advantages over ConvNets that make them highly valuable for ecologists and evolutionary biologists. Contrary to ConvNets, which require a huge amount of data for training (i.e. to learn filter selectivities), S2 filters in HMAX can be learned from a few images, and possibly a single image. Moreover, contrary to ConvNets, with HMAX it is straightforward to visualize the selectivity of neurons (i.e. filters) and to analyse which neurons are activated and which are not. This is convenient, e.g., for revealing those features that most influence the similarity between two phenotypes. HMAX models thus have high explanatory power, which ConvNets still critically lack.

Compared to the classic HMAX, sparse-HMAX showed reduced performance both for estimating facial similarity in mandrills and in performance tests (see SI). However, our goal in proposing a sparse implementation of HMAX was not to maximize performance but to make the model biologically more realistic. Furthermore, *sparse*-HMAX provides a tool for estimating neuronal sparseness and thus efficiency in information processing (Renoult & Mendelson 2019). A growing body of psychological studies suggests that, when given a choice, humans tend to prefer stimuli that are efficiently processed by the brain (Winkielman *et al.* 2003; Reber, Schwarz & Winkielman 2004; Redies 2007). This finding is appealing for the field of evolutionary biology as it could shed light on the mechanisms underlying the evolution of complex and extravagant communication signals in animals (Renoult & Mendelson 2019). Studying the efficiency of information processing outside laboratories of neurophysiology and in non-model animal species, however, will require models such as sparse-HMAX that quantify processing efficiency in animal brains. Visual ecology and evolutionary biology are only beginning to embrace the benefits of methods developed in computer vision and computational neuroscience. One aim of this article was to make one step forward toward a better connection between these fields of research.

## Acknowledgements

The authors thank Alexandre Faucher for preliminary studies on sparseness and HMAX. This study is funded by National Science Foundation grant IOS-1708543, and by the LABEX NUMEV, AGRO and CEMEB from University of Montpellier.

## Authors’ contributions

JPR and TM designed the study and wrote the first draft of the article; JPR performed the facial similarity analyses; BG wrote the Matlab program including its documentation, made performance tests and is managing the GitHub repository; JD, FG and FM developed the sparse-HMAX algorithm; AP performed the landmark analysis; all authors contributed to the final version of the manuscript.

## Data repository

MATLAB codes and documentation available at https://github.com/EEVCOM-Montpellier/HMAX

## References

Abbas, F. & Meyer, M.P. (2014) Fish vision: Size selectivity in the zebrafish retinotectal pathway. Current Biology, 24, R1048–1050.

Anselmi, F., Leibo, J.Z., Rosasco, L., Mutch, J., Tacchetti, A. & Poggio, T. (2016) Unsupervised learning of invariant representations. Theoretical Computer Science, 633, 112–121.

Attwell, D. & Laughlin, S.B. (2001) An energy budget for signaling in the grey matter of the brain. Journal of Cerebral Blood Flow & Metabolism, 21, 1133–1145.

Ben-Tov, M., Kopilevich, I., Donchin, O., Ben-Shahar, O., Giladi, C. & Segev, R. (2013) Visual receptive field properties of cells in the optic tectum of the archer fish. Journal of Neurophysiology, 110, 748–759.

Bower, S., Suomi, S.J. & Paukner, A. (2012) Evidence for kinship information contained in the rhesus macaque (*Macaca mulatta*) face. Journal of Comparative Psychology, 126, 318–323.

Brincat, S.L. & Connor, C.E. (2004) Underlying principles of visual shape selectivity in posterior inferotemporal cortex. Nature Neuroscience, 7, 880–886.

Carlson, E.T., Rasquinha, R., Zhang, K. & Connor, C.E. (2011) A sparse object coding scheme in area V4. Current Biology, 21, 288–293.

Caves, E.M., Brandley, N.C. & Johnsen, S. (2018) Visual acuity and the evolution of signals. Trends in Ecology & Evolution, 33, 358–372.

Cummings, M.E. & Endler, J.A. (2018) 25 Years of Sensory Drive: The evidence and its watery bias. Current Zoology, 64, 471–484.

Dal Martello, M.F. & Maloney, L.T. (2006) Where are kin recognition signals in the human face? Journal of Vision, 6, 2–2.

Daw, N.W. (1968) Colour – coded ganglion cells in the goldfish retina: extension of their receptive fields by means of new stimuli. The Journal of Physiology, 197, 567–592.

Derrington, A.M., Krauskopf, J. & Lennie, P. (1984) Chromatic mechanisms in lateral geniculate nucleus of macaque. The Journal of Physiology, 357, 241–265.

DiCarlo, J.J., Zoccolan, D. & Rust, N.C. (2012) How does the brain solve visual object recognition? Neuron, 73, 415–434.

Endler, J.A., Westcott, D.A., Madden, J.R. & Robson, T. (2005) Animal visual systems and the evolution of color patterns: Sensory processing illuminates signal evolution. Evolution, 59, 1795–1818.

Engelage, J. & Bischof, H. (1996) Single-cell responses in the ectostriatum of the zebra finch. Journal of Comparative Physiology A, 179, 785–795.

Felleman, D.J. & Van Essen, D.C. (1991) Distributed hierarchical processing in the primate cerebral cortex. Cerebral Cortex, 1, 1–47.

Ferster, D. & Miller, K.D. (2000) Neural mechanisms of orientation selectivity in the visual cortex. Annual Review of Neuroscience, 23, 441–471.

Freedman, D.J., Riesenhuber, M., Poggio, T. & Miller, E.K. (2005) Experience-dependent sharpening of visual shape selectivity in inferior temporal cortex. Cerebral Cortex, 16, 1631–1644.

Frost, B.J., Scilley, P.L. & Wong, S.C.P. (1981) Moving background patterns reveal double-opponency of directionally specific pigeon tectal neurons. Experimental Brain Research, 43, 173–185.

Gegenfurtner, K.R. (2003) Cortical mechanisms of colour vision. Nature Reviews Neuroscience, 4, 563–572.

Graham, D.J. & Field, D.J. (2006) Sparse coding in the neocortex. Evolution of Nervous Systems, 3, 181–187.

Heeger, D.J. (1992) Normalization of cell responses in cat striate cortex. Visual Neuroscience, 9, 181–197.

Hromadka, T., DeWeese, M.R. & Zador, A.M. (2008) Sparse representation of sounds in the unanesthetized auditory cortex. PLoS Biology, 6, e16.

Hu, X., Zhang, J., Li, J. & Zhang, B. (2014) Sparsity-regularized HMAX for visual recognition. PloS One, 9, e81813.

Hubel, D.H. & Wiesel, T.N. (1962) Receptive fields, binocular interaction and functional architecture in the cat’s visual cortex. The Journal of Physiology, 160, 106–154.

Johnson, E.N., Hawken, M.J. & Shapley, R. (2001) The spatial transformation of color in the primary visual cortex of the macaque monkey. Nature Neuroscience, 4, 409–416.

Johnson, M.H., Dziurawiec, S., Ellis, H. & Morton, J. (1991) Newborns’ preferential tracking of face-like stimuli and its subsequent decline. Cognition, 40, 1–19.

Jones, J.P. & Palmer, L.A. (1987) An evaluation of the two-dimensional Gabor filter model of simple receptive fields in cat striate cortex. Journal of Neurophysiology, 58, 1233–1258.

Kelber, A. (2002) Pattern discrimination in a hawkmoth: Innate preferences, learning performance and ecology. Proceedings of the Royal Society B, 269, 2573–2577.

Kriegeskorte, N. (2015) Deep neural networks: A new framework for modeling biological vision and brain information processing. Annual Review of Vision Science, 1, 417–446.

LeCun, Y., Bengio, Y. & Hinton, G. (2015) Deep learning. Nature, 521, 436.

Lee, H., Battle, A., Raina, R. & Ng, A.Y. (2007) Efficient sparse coding algorithms. Advances in Neural Information Processing Systems, 19, 801–808.

Lennie, P. (2003) The cost of cortical computation. Current Biology, 13, 493–497.

Li, D.-P., Xiao, Q. & Wang, S.-R. (2006) Feedforward construction of the receptive field and orientation selectivity of visual neurons in the pigeon. Cerebral Cortex, 17, 885–893.

Mély, D.A. & Serre, T. (2017) Towards a theory of computation in the visual cortex. Computational and Cognitive Neuroscience of Vision (ed. Zhao, Q), pp. 59–84. Springer.

Mutch, J. & Lowe, D.G. (2008) Object class recognition and localization using sparse features with limited receptive fields. International Journal of Computer Vision, 80, 45–57.

Ng, A.Y. (2004) Feature selection, L1 vs. L2 regularization, and rotational invariance. Proceedings of the Twenty-First International Conference on Machine learning, pp. 78. ACM.

Olshausen, B.A. & Field, D. (1996) Emergence of simple-cell receptive field properties by learning a sparse code for natural images. Nature, 381, 607–609.

Olshausen, B.A. & Field, D.J. (1997) Sparse coding with an overcomplete basis set: A strategy employed by V1? Vision Research, 37, 3311–3325.

Olshausen, B.A. & Field, D.J. (2004) Sparse coding of sensory inputs. Current Opinion in Neurobiology, 14, 481–487.

Pitkow, X. & Meister, M. (2012) Decorrelation and efficient coding by retinal ganglion cells. Nature Neuroscience, 15, 628–635.

Poggio, T. & Serre, T. (2013) Models of visual cortex. Scholarpedia, 8, 3516.

Reber, R., Schwarz, N. & Winkielman, P. (2004) Processing fluency and aesthetic pleasure: is beauty in the perceiver’s processing experience? Personality and Social Psychology Review, 8, 364–382.

Redies, C. (2007) A universal model of esthetic perception based on the sensory coding of natural stimuli. Spatial Vision, 21, 97–117.

Reid, R.C. & Usrey, W.M. (2004) Functional connectivity in the pathway from retina to striate cortex. The visual Neurosciences, 1, 673–679.

Renoult, J.P., Kelber, A. & Schaefer, H.M. (2017) Colour spaces in ecology and evolutionary biology. Biological Reviews, 92, 292–315.

Renoult, J.P. & Mendelson, T.C. (2019) Processing bias: Extending sensory drive to include efficacy and efficiency in information processing. arXiv Preprints, arXvive: 1901.00782

Riesenhuber, M. & Poggio, T. (1999) Hierarchical models of object recognition in cortex. Nature Neuroscience, 2, 1019–1025.

Scherer, D., Môller, A. & Behnke, S. (2010) Evaluation of pooling operations in convolutional architectures for object recognition. Artificial Neural Networks-ICANN 2010, 92–101.

Sengpiel, F. & Kind, P.C. (2002) The role of activity in development of the visual system. Current Biology, 12, R818–R826.

Sengpiel, F., Stawinski, P. & Bonhoeffer, T. (1999) Influence of experience on orientation maps in cat visual cortex. Nature Neuroscience, 2.

Serre, T. (2013) Hierarchical models of the visual system. Encyclopedia of Computational Neuroscience, 1–12.

Serre, T., Kouh, M., Cadieu, C., Knoblich, U., Kreiman, G. & Poggio, T. (2005) A theory of object recognition: computations and circuits in the feedforward path of the ventral stream in primate visual cortex. Massachusetts Institute of Technology, Cambridge, MA Center for Computational Learning.

Serre, T. & Riesenhuber, M. (2004) Realistic modeling of simple and complex cell tuning in the HMAX model, and implications for invariant object recognition in cortex. DTIC Document.

Shapley, R. & Hawken, M.J. (2011) Color in the cortex: single-and double-opponent cells. Vision Research, 51, 701–717.

Smith, A.T., Singh, K.D., Williams, A.L. & Greenlee, M.W. (2001) Estimating receptive field size from fMRI data in human striate and extrastriate visual cortex. Cerebral Cortex, 11, 1182–1190.

Soto, F.A. & Wasserman, E. (2012) Visual object categorization in birds and primates: Integrating behavioral, neurobiological, and computational evidence within a “general process” framework. Cognitive, Affective, & Behavioral Neuroscience, 12, 220–240.

Stoddard, M.C., Kilner, R.M. & Town, C. (2014) Pattern recognition algorithm reveals how birds evolve individual egg pattern signatures. Nature communications, 5, 4117.

Theriault, C., Thome, N. & Cord, M. (2013) Extended coding and pooling in the hmax model. IEEE Transactions on Image Processing, 22, 764–777.

Thorpe, S., Fize, D. & Marlot, C. (1996) Speed of processing in the human visual system. Nature, 381, 520–522.

Tootell, R.B., Switkes, E., Silverman, M.S. & Hamilton, S. (1988) Functional anatomy of macaque striate cortex. II. Retinotopic organization. Journal of Neuroscience, 8, 1531–1568.

Treue, S. (2003) Visual attention: The where, what, how and why of saliency. Current Opinion in Neurobiology, 13, 428–432.

VanRullen, R. & Koch, C. (2003) Visual selective behavior can be triggered by a feedforward process. Journal of Cognitive Neuroscience, 15, 209–217.

Winkielman, P., Schwarz, N., Fazendeiro, T. & Reber, R. (2003) The hedonic marking of processing fluency: Implications for evaluative judgment. The psychology of evaluation: Affective processes in cognition and emotion (eds J. Musch & K.C. Klauer), pp. 189–217. Psychology Press.

Zeki, S., Watson, J.D., Lueck, C.J., Friston, K.J., Kennard, C. & Frackowiak, R. (1991) A direct demonstration of functional specialization in human visual cortex. Journal of Neuroscience, 11, 641–649.

Zhang, J., Barhomi, Y. & Serre, T. (2012) A new biologically inspired color image descriptor. Computer vision-ECCV 2012, 312–324.

