## Supplementary Material for "Modelling the Perception of Colour Patterns in Vertebrates with HMAX"

**Supporting Information**

### Abbreviations

| | |
|---|---|
| $b$ | index of a photoreceptor channel |
| $C$i | complex-cells layer i |
| $DO$ | double-opponent cells |
| $h$ | height, in pixel, of the input matrix |
| $j$ | index of a colour channel |
| $k$ | length of $\vec{\xi}$ |
| $k'$ | scale parameter in $SO$ normalisation |
| $I$ | input matrix of layer $S1$ or $SO$ |
| $I_b$ | photoreceptor excitation channel |
| IT | infero-temporal cortex |
| $\vec{f}_n^{S2}$ | a filter generated by sparse coding converted to column-vector |
| $F^{Si}/F^{Di}$ | a filter of $S$i/$D$i |
| $\mathbf{F}^{Si}/\mathbf{F}^{Di}$ | set of filters of $S$i/$D$i |
| $L$ | long-wavelength sensitive photoreceptor |
| LGN | lateral geniculate nucleus |

| | | |
|---|---|---|
| 25 | $M^{Si/Di}$ | a feature map of $S$i/$D$i |
| 26 | $\mathbf{M}^{Si/Di}$ | set of feature maps of $S$i/$D$i |
| 27 | $M$ | medium-wavelength sensitive photoreceptor |
| 28 | $m$ | number of distinct non-overlapping patches in $M^{C1}_{\theta,\sigma}$ |
| 29 | $\vec{M}^{C2}$ | output of Hmax, typically a vector of approx. 1000 values |
| 30 | $n$ | index of filter $S2$, and within vector $\vec{M}^{C2}$ |
| 31 | $N$ | number filters in $S2$, and length of $\vec{M}^{C2}$ |
| 32 | $nch$ | number of channels in $SO$ (including luminance channel) |
| 33 | $npe$ | number of photoreceptor excitation maps |
| 34 | $P$ | patch = a small area of $M^{C1}_{\sigma}$ |
| 35 | $q$ | size of a Gabor filter = RF of neurones in V1 (in mammals) |
| 36 | $q'$ | size of a $S2$ filter = RF of neurons in V4/IT (in mammals) |
| 37 | $r$ | size of max pooling filters |
| 38 | $Re$ | response of a filter to a patch $P$ from $M^{C1}_{\sigma}$ |
| 39 | RF | receptive field of a neuron |
| 40 | $s$ | scaling parameter in response function $R$ |
| 41 | $S$ | short-wavelength sensitive photoreceptor |
| 42 | $\mathbf{S}$ | matrix of filter weights in the sparse coding minimization function |
| 43 | SVM | Support Vector Machine |
| 44 | $sc$ | number of different scales for Gabor filters |
| 45 | $S$i | simple-cells layer i |
| 46 | $SO$ | single-opponent cells |
| 47 | $t$ | number of different Gabor filter orientations in $\mathbf{F}^{S1}$ and $\mathbf{F}^{DO}$ |
| 48 | $t^{SO}$ | number of different Gabor filter orientations in $\mathbf{F}^{SO}$ |
| 49 | $u$ | number of different Gabor filter sizes |

| | |
|---|---|
| $u'$ | number of different $S2$ filters sizes |
| Vi | cortical visual area i |
| $w$ | width, in pixel, of the input matrix |
| $(x,y)$ | spatial coordinates within a feature map |
| (x,y) | spatial coordinates of a filter centre |
| $x_g$,$y_g$ | spatial position within a Gabor feature map of a filter centre |
| **X** | matrix concatenating all $\vec{\xi}$ |
| $\alpha$ | normalisation factor in response function $R$ |
| $\beta$ | regularisation parameter of the sparse coding minimization function |
| $\gamma$ | aspect ratio of a Gabor filter |
| $\vec{\xi}$ | a patch of $M_{\sigma,\theta}^{C1}$ transformed to column-vector |
| $\lambda$ | wavelength parameter in Gabor function |
| $\sigma$ | scale of Gabor filter |
| $\sigma'$ | semi-saturation parameter in SO normalisation |
| σ | one random index of within $\vec{\Sigma}$ |
| $\theta$ | orientation of a Gabor filter in $\mathbf{F}^{S1}$ and $\mathbf{F}^{DO}$ |
| $\theta^{SO}$ | orientation of a Gabor filter in $\mathbf{F}^{SO}$ |
| * | polarity of a Gabor filter (modelling $SO$ cells) |
| $\mathbf{\Omega}$ | matrix of weights in the opponent and luminance function of $SO$ |

**Performance in object recognition**

We tested the performance of our versions of (colour) HMAX and (colour) *sparse*-HMAX in a recognition task using the Caltech101 object dataset (Fei-Fei, Fergus & Perona 2007). This dataset represents 102 categories, each one containing a minimum of 31 pictures. Analyses were performed with the 102 categories, and with a reduced dataset of only 20 categories. Images were resized to 140 pix for the smallest side, keeping the aspect ratio unchanged (Table S1). We used 15 images per category to first learn the filter set $\mathbf{F}^{S2}$, extracting 1,000 filters from $C1$ with HMAX and 4,000 filters with colour HMAX (1,000 for each *DO* channel), and learning 250 filters with *sparse*-HMAX (both greyscale and colour versions). We used a red-cyan opponent channel in addition to the yellow-blue, red-green and achromatic channels (Zhang, Barhomi & Serre 2012). For the classification task, we randomly selected 31 pictures from each of the 102 categories and encoded them with the previously learned HMAX models (for parameter details, see Table S1). Among these 31 pictures, 25 were used for training the classifier and six for testing. The multiclass classifier consisted of 102 binary linear Support Vector Machines (one per category). During training, we applied a cross-validation procedure with five clusters to limit over-fitting.

**Table S1.** HMAX parameters (see documentation of the MATLAB package) used to learn $F^{S2}$ filters for testing performances with Caltech101.

| **General parameters** | |
|---|---|
| Image size | 140 px |
| Gabor - Nb orientations | 4 |
| Gabor - Aspect ratio | 0.3 |
| Gabor - Effective width | [2.8; 3.6; 4.5; 5.4; 6.3; 7.3; 8.2; 9.2; 10.2; 11.3; 12.3; 13.4; 14.6; 15.8; 17; 18.2 ] |
| Gabor - Wavelength | [3.5; 4.6; 5.6; 6.8; 7.9; 9.1; 10.3; 11.5; 12.7; 14.1; 15.4; 16.8; 18.2; 19.7; 21.2; 22.8] |
| Gabor - Size | [7; 9; 11; 13; 15; 17; 19; 21; 23; 25; 27; 29; 31; 33; 35; 37] |
| **HMAX and colour HMAX** | |
| MaxPoolingSizes | [8; 10; 10; 12; 12; 14; 14; 16; 16; 18; 18; 20; 20; 22; 22] |
| Number of filters | 1000 |
| **Colour (sparse) HMAX** | |

| | |
|---|---|
| Number of image channels | 4 |
| **$F^{S2}$ in sparse HMAX** | |
| Size | 12 |
| NbPatches | 10000 |
| BatchSize | 1000 |
| NbIterations | 3 |
| Penalty | 0.4 |
| NbFilters | 250 |

Our version of HMAX yielded similar performance to that of Serre *et al.* (2007) with Caltech20 (c.a. 58% of correct classification) and slightly better performance with Caltech101 (c.a. 32% versus 27%), but was c.a. 25% faster (see the "Result" section in the documentation of our MATLAB code at: https://eevcom-montpellier.github.io/hmax-documentation/methodology-and-results/). With colour HMAX, our version performed markedly better compared to that of Zhang, Barhomi & Serre (2012) with Caltech20 (64% versus 51%) and slightly better with Caltech101 (30% versus 26%) although it was between 17% and 25% slower. *sparse*-HMAX showed lower performance compared to HMAX in all analyses (e.g., 46% and 54% for greyscale and colour versions, respectively, with Caltech20).

### Mandrill face similarity

#### *Image acquisition*

Pictures of mandrill faces were taken on anesthetized animals that were all individually recognized and during different trapping sessions conducted between 2012 and 2015 in Gabon (for details, see Beaulieu *et al.* 2017). Some individuals were trapped and photographed only once, while others could be photographed several times over one, two or three consecutive years.

*Landmarks*

We positioned twenty-four landmarks on mandrills' faces (see: Bower, Suomi & Paukner 2012), avoiding chin and mouth because of differences in head positions between images (Figure S1). We used tpsDig2© 2.22 software to position landmarks and then imported the 2D Cartesian landmark coordinates (.TPS files) into MorphoJ v.1.06d software in order to perform a full Procrustes superimposition (as in Webster & Sheets 2010), that is, a shape transformation that applies scaling, translation and rotation to the imported sets of landmarks. To do so, we aligned landmarks with the x-axis of the reference base coordinate system, using landmarks 9 and 13 (Table S2). We then used R v3.03 software in order to estimate visual dissimilarity across pairs of pictures based on Procrustes coordinates. We calculated a morphometric distance (*d*) for each dyad of pictures as follow:

$$dxy = \sqrt{\sum_{i,j}^{k,p} (Xij - Yij)^2}$$

where *x*, *y* and *k* are the dimensions of the coordinate system (here, $k$=2) and *p* the number of landmarks (here, $p$=24). *X* and *Y* represents the Procrustes coordinates of the pictures *x* and *y* (respectively), *i* and *j* correspond to the horizontal and vertical dimensions of the coordinate system. Morphometric distances are positive; an elevated distance indicates dissimilar faces.

**Table S2. Position of landmarks.**

| Landmark | Position on face |
|---|---|
| 1 | Left extremity of the left eye |
| 2 | Right extremity of the left eye |
| 3 | Left extremity of the right eye |
| 4 | Right extremity of the right eye |
| 5 | Superior point of the left eye brow arch |
| 6 | Inferior point of the left orbit |
| 7 | Superior point of the right eye brow arch |
| 8 | Inferior point of the right orbit |
| 9 | Left point of the left orbit |
| 10 | Up right corner of the left orbit, where the nose bridge starts |
| 11 | Central point of the highest wrinkle between the two eyes |
| 12 | Up left corner of the right orbit, where the nose bridge starts |
| 13 | Right point of the right orbit |
| 14 | Up central point of the suborbital bulge |
| 15 | Left point of the top of the nose, at the end of the nose bridge |
| 16 | Right point of the top of the nose, at the end of the nose bridge |
| 17 | Tip of the nose, where nasal cartilage of both sides of the nose joined at the center |
| 18 | Left extremity of the bottom of the left nostril |
| 19 | Right extremity of the bottom of the right nostril |
| 20 | Left point at the bottom of the nasal septum |
| 21 | Right point at the bottom of the nasal septum |

| | |
|---|---|
| 22 | Central point of the upper lip, perpendicular to the nose axis |
| 23 | Highest point of the left nostril |
| 24 | Highest point of the right nostril |

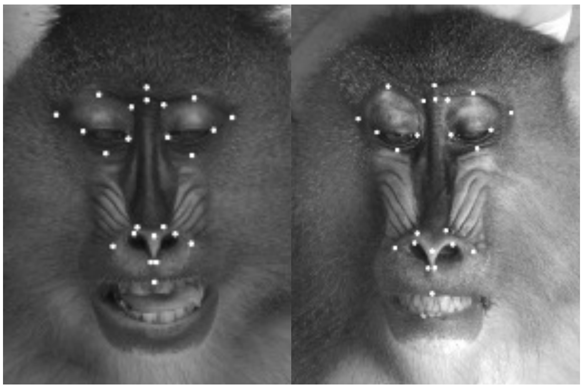

**Figure S1. Example of images used for landmark analyses.**

*Image pre-processing for computer vision analyses*

Images were first converted to greyscale and contrast adjusted by histogram equalization. Head fur was replaced by a black background and a fade transition was added manually between the bare face and the background. Portrait images were squared by horizontal padding and resized to 400 x 400 px.

*SURF*

Speeded Up Robust Features (SURF; Bay *et al.* 2008) is an algorithm for automatically detecting rotation and scale invariant features in an image. It is very close to the Scale Invariant Feature Transform algorithm (SIFT; Lowe 2004), another feature detector widely used in computer vision, but it is generally faster (Bay *et al.* 2008).

We estimated the similarity between mandrill faces based on the number of homologous features shared between two images, using an approach originally developed with SIFT and presented in details in Stoddard, Kilner & Town (2014). The algorithm first detects key points based on the determinant of the Hessian matrix, and then extracts features around each key point based on the summed response to Haar wavelets. Then, the algorithm identifies homologous features in two images by calculating the pairwise distance (i.e., sum of square differences) between all features and by comparing these distances to a “distance threshold”. The distance threshold represents a percent of the distance to a perfect match: two features are homologous if their distance is lower than the threshold. In addition, ambiguous matches are rejected based on two different approaches: when the ratio between the distance-to-the-best-match and the distance-to-the-second-best-match is greater than a “ratio-threshold”; and by identifying outlying matches with the Random Sample Consensus (RANSAC) algorithm. A score of similarity corresponds to the number of non-ambiguous matches divided by the number of maximum possible matches. For two images $I$ and $J$, the final similarity score is conservatively calculated as the lowest of the two scores calculated when image $I$ is compared to image $J$, and when image $J$ is compared to image $I$. SURF analyses were performed with the Computer Vision Toolbox of MATLAB. Different combinations of parameter values have been tested; Table S3 provides settings that yielded the best performance (i.e., rank sum; see main text).

**Table S3**. Selected parameters for estimating facial similarity in mandrills using the SURF algorithm.

| Task | Parameter | Matlab nomenclature | Value |
|---|---|---|---|
| Keypoint detection and feature extraction | Strongest feature threshold | MetricThreshold | 3 |
| | Number of octaves | NumOctaves | 3 |
| | Number of scales per octave | NumScaleLevels | 4 |
| | Feature size | SURFSize | 128 |
| Feature matching | Distance threshold | MatchThreshold | 3 |
| | Ratio threshold | MaxRatio | 0.7 |

*HMAX*

We first learned $\mathbf{F}^{S2}$ filters by training HMAX to encode a dataset of 132 images of pre-processed as described previously. The individuals (n=76) displayed on this learning dataset were different from those included in the distance measurement analysis. Then, we used this pretrained HMAX algorithm to encode the same pictures of mandrill faces as in the landmark and SURF analyses (“testing” in Table S2). Pairwise distances between images were then calculated as one minus the cosine of the included angle between $C2$ vectors. We used the same parameter settings as in Caltech101 analysis, except that images were larger.

*Score comparison*

Because similarity metrics differ between methods (landmark, SURF and HMAX), we compared methods using a rank-based approach. Similarity scores calculated for all pairs of images were ranked in ascending order, and then we calculated the sum of ranks of pairs corresponding to same-individual images. The rank sums for landmark, SURF and HMAX were compared to a null distribution obtained after 9999 permutations of ranks, and to an ideal observer attributing the highest ranks (i.e. highest similarity, or smallest distances) to all same-individual pairs.

### References


Bay, H., Ess, A., Tuytelaars, T. & Van Gool, L. (2008) Speeded-up robust features (SURF). *Computer Vision and Image Understanding,* **110,** 346-359.

Beaulieu, M., Benoit, L., Abaga, S., Kappeler, P.M. & Charpentier, M.J.E. (2017) Mind the cell: Seasonal variation in telomere length mirrors changes in leucocyte profile. *Molecular Ecology,* **26,** 5603-5613.

Bower, S., Suomi, S.J. & Paukner, A. (2012) Evidence for kinship information contained in the rhesus macaque (*Macaca mulatta*) face. *Journal of Comparative Psychology,* **126,** 318-323.

Fei-Fei, L., Fergus, R. & Perona, P. (2007) Learning generative visual models from few training examples: An incremental bayesian approach tested on 101 object categories. *Computer Vision and Image Understanding,* **106,** 59-70.

Lowe, D.G. (2004) Distinctive image features from scale-invariant keypoints. *International Journal of Computer Vision,* **60,** 91-110.

Serre, T., Wolf, L., Bileschi, S., Riesenhuber, M. & Poggio, T. (2007) Robust object recognition with cortex-like mechanisms. *IEEE Transactions on Pattern Analysis & Machine Intelligence*, 411-426.

Stoddard, M.C., Kilner, R.M. & Town, C. (2014) Pattern recognition algorithm reveals how birds evolve individual egg pattern signatures. *Nature Communications,* **5,** 4117.

Webster, M. & Sheets, H.D. (2010) A practical introduction to landmark-based geometric morphometrics. *The Paleontological Society Papers,* **16,** 163-188.

Zhang, J., Barhomi, Y. & Serre, T. (2012) A new biologically inspired color image descriptor. *Computer Vision–ECCV 2012*, 312-324.
